# The *Brachypodium distachyon* cold-acclimated plasma membrane proteome is primed for stress resistance

**DOI:** 10.1101/2021.04.23.441164

**Authors:** Collin L. Juurakko, Melissa Bredow, Takato Nakayama, Hiroyuki Imai, Yukio Kawamura, George C. diCenzo, Matsuo Uemura, Virginia K. Walker

## Abstract

In order to survive sub-zero temperatures, some plants undergo cold acclimation where low, non-freezing temperatures and/or shortened day lengths allow cold hardening and survival during subsequent freeze events. Central to this response is the plasma membrane, where low-temperature is perceived and cellular homeostasis must be preserved by maintaining membrane integrity. Here, we present the first plasma membrane proteome of cold-acclimated *Brachypodium distachyon*, a model species for the study of monocot crops. A time course experiment investigated cold acclimation-induced changes in the proteome following two-phase partitioning plasma membrane enrichment and label-free quantification by nano-liquid chromatography mass spectrophotometry. Two days of cold acclimation were sufficient for membrane protection as well as an initial increase in sugar levels, and coincided with a significant change in the abundance of 154 proteins. Prolonged cold acclimation resulted in further increases in soluble sugars and abundance changes in more than 680 proteins, suggesting both a necessary early response to low-temperature treatment, as well as a sustained cold acclimation response elicited over several days. A meta-analysis revealed that the identified plasma membrane proteins have known roles in low-temperature tolerance, metabolism, transport, and pathogen defense as well as drought, osmotic stress and salt resistance suggesting crosstalk between stress responses, such that cold acclimation may prime plants for other abiotic and biotic stresses. The plasma membrane proteins identified here present keys to an understanding of cold tolerance in monocot crops and the hope of addressing economic losses associated with modern climate-mediated increases in frost events.

## INTRODUCTION

Changing climatic conditions are associated with unpredictable weather patterns that can have devastating consequences on crop success (Raza *et al*. 2019). Higher average temperatures and an increased frequency of winter freeze-thaw events present major challenges in temperate regions and can result in delayed bud-burst and freeze-induced injury, with acute exposure to temperatures below a thermal optimum generating chilling stress (Aroca *et al*. 2012; Tedla *et al*. 2020). Low-temperature effects include lower rates of biochemical and metabolic reactions, a loss in membrane fluidity, increased water viscosity, decreased water uptake in roots, attenuated activity of numerous proteins and enzymes, as well as delayed energy dissipation associated with reduced photosynthesis and cellular respiration (Aroca *et al*. 2012). As temperatures lower further, there is the added challenge of freezing stress, resulting from the growth of extracellular ice crystals that physically damage plasma membranes (PMs), as well as cellular dehydration, which in turn results in the generation of reactive oxygens that together can ultimately lead to the collapse of membrane structures (Pearce 2001). Given the importance of cereal crops for food security, it is crucial to identify proteins associated with freeze protection in any effort to improve freezing resistance.

Prior to anthropogenic-induced climate change, plants in their native range would presumably only rarely be exposed to atypically acute and thus fatal exposure to freezing. Instead, cold acclimation (CA) - a process induced by low, non-freezing temperatures and/or shortened day lengths and involving epigenetic, biochemical, metabolic and physiological changes - results in cold-hardening (Thomashow 1999, 2010; Fürtauer *et al*. 2019). Cold-hardened plants often accumulate unsaturated fatty acids, produce cryoprotective metabolites including soluble sugars and amino acids to mitigate osmotic stress, and elevated expression of molecular chaperons and reactive oxygen species scavengers (Suzuki and Mittler 2006). Many plants undergo CA, but the degree of their subsequent freezing tolerance is primarily determined by geographic origin, with some cold-hardy plants withstanding temperatures as low as -30 °C (Lee *et al*. 2012; Zuther *et al*. 2012; Colton-Gagnon *et al*. 2014). Many of these species upregulate the expression of cryoprotective proteins including cold-responsive (COR) proteins, dehydrins, and ice-binding proteins (Liu *et al*. 2017; Bredow and Walker 2017).

Low-temperature sensing is initiated at the PM, in part by increased membrane rigidity and activation of mechanosensitive Ca^2+^ channels that initiate downstream signaling events key to cold tolerance (Mori *et al*. 2018; Yuan *et al*. 2018). In rice, *Oryza sativa,* low-temperature is perceived by a PM G-protein signaling receptor, COLD1, required for Ca^2+^ channel activation (Ma *et al*. 2015). Analyses of the PM-associated proteome of thale cress, *Arabidopsis thaliana,* hereinafter *Arabidopsis* (Kawamura and Uemura 2002; Miki *et al*. 2019; Li *et al*. 2020), winter rye, *Secale cereale* (Uemura and Yoshida 1984), and oat, *Avena sativa* (Takahashi *et al*. 2010) suggest that the PM is also important for initiating cold-induced changes in these plants. Once cold-induced signaling has commenced, there is an accumulation of membrane-stabilizing COR proteins, a remodelling of the PM, as well as an upregulation of osmolyte synthases that help protect against dehydration-induced cell lysis (Minami *et al*. 2009; Takahashi *et al*. 2018).

Although not a crop plant, purple false brome *Brachypodium distachyon* (hereinafter, *Brachypodium*) is a model monocot with physiological similarity and high synteny with agriculturally important grasses such as rice and wheats (Triticeae). Evidence shows that *Brachypodium* evolved in Mediterranean regions and some cultivars tolerate low-temperatures (Ryu *et al*. 2014; Colton-Gagnon *et al*. 2014; Bredow *et al*. 2016). Following CA, these survive to -10 °C (Colton-Gagnon *et al*. 2014) with a diurnal freezing treatment allowing tolerance down to -12 °C (Mayer *et al*. 2020). Here, we examine the profile of compatible solutes accumulated after CA, quantify cold-induced membrane protection, and present the first proteomic analysis of cold-acclimated (CA) *Brachypodium* from PM-enriched microsomal fractions in order to further advance our understanding of freeze-tolerance and the low-temperature protection in this model monocot.

## MATERIALS AND METHODS

### Plant growth and maintenance

*Brachypodium* seeds (ecotype: *Bd*21) were sown in potting soil and grown in a temperature controlled chamber on a 20 h light (∼100 µmol m^-2^ s^-1^; 22 °C) and 4 h dark (22 °C) light cycle. CA *Brachypodium* were grown under standard conditions for three weeks and transferred to a low-temperature chamber (2 °C, 12 h light as indicated above; 12 h dark), for two to eight days with experimental groupings for two, four, six, and eight days designated CA2, CA4, CA6 and CA8, respectively.

Non-acclimated (NA) *Brachypodium* were three-weeks-old at the time of use and not incubated further.

### Electrolyte leakage

In order to assess the level of membrane damage associated with freezing, electrolyte leakage assays were conducted using NA or CA plants as described previously (Bredow *et al*. 2016). Briefly, one leaf was cut from the base of each plant, placed in 100 μL of deionized water, and immersed in a programmable circulating ethylene glycol bath set to 0 °C. After lowering the temperature to -1 °C over 30 min, the sample was nucleated with a single deionized ice chip to initiate ice crystal growth. The temperature of the bath was then lowered by 1 °C every 15 min to a final temperature of -2 to -10 °C. Samples were allowed to recover overnight at 4 °C, transferred into conical tubes containing 25 mL of deionized water, and then gently agitated at 150 rpm for 18 h. Conductivity measurements were taken before (C_i_) and after (C_f_) autoclaving samples to account for the total leaf mass, using a direct reading conductivity meter (TwinCond, Horiba, Kyoto, Japan) and presented as a percentage (100 * C_i_ / C_f_) with 10 individual plants for each independent line, with the procedure carried out with three biological replicates.

### Compatible solute profiling

Leaves (∼400 mg fresh weight) were frozen in liquid nitrogen and ground with a plastic pestle in 0.25 mL of 80% ethanol with fucose as an internal standard.

Homogenates were transferred into 1.5 mL microcentrifuge tubes and incubated at 80 °C for 30 min with occasional mixing. Samples were then centrifuged at 12,000 x *g* for 5 min at room temperature, the supernatants collected and the pellets re-extracted twice with 80% ethanol at 80 °C and then centrifuged as described above. The supernatants were combined and dried under a dried N_2_ gas stream at 80 °C. These extracts were then dissolved in methanol, followed by the addition of chloroform and water (a final ratio of chloroform:methanol:water, 1:1:0.9, v/v/v) to remove leaf pigments. After centrifugation at 600 x *g* for 5 min, the upper aqueous phase was collected and dried as described above. Dried samples were dissolved in water, centrifuged at 12,000 x *g* for 5 min, and the supernatants filtered through a 0.2 μm membrane filter and subsequently analyzed by high pressure liquid chromatography (HPLC) using a tandem-connected Sugar KS801 and KS802 column (Shodex, Tokyo, Japan) at 80 °C with a refraction index detector. The samples (50 μL) were injected and eluted with ultrapure water at a flow rate of 0.4 mL min^-1^. Each sugar was identified by comparison of its retention time relative to that of authentic standard sugars and quantified as a ratio of the area of the sample peak relative to that of the internal standard. All assays were replicated four times.

### Microsome isolation and plasma membrane enrichment

The extraction of microsomal fractions from plant tissue and subsequent enrichment of PMs was performed as detailed previously (Kamal *et al*. 2020). Briefly, ∼30 g of leaf tissue was cut into small pieces in 300 mL of pre-chilled homogenizing medium (0.5 M sorbitol, 50 mM MOPS-KOH (pH 7.6), 5 mM EGTA (pH 8.0), 5 mM EDTA (pH 8.0), 1.5% (w/v) PVP-40, 0.5% (w/v) BSA, 2.5 mM PMSF, 4 mM SHAM, 2.5 mM DTT). Tissue was further homogenized using a Polytron generator (PT10SK, Kinematica Inc., Lucerne, Switzerland) and filtrates were collected by sieving the homogenate through cheesecloth and removing cellular debris by centrifugation (10,000 x *g* for 10 min). Microsomal membrane fractions were collected by centrifugation at 231,000 x *g* for 35 min and homogenized in 1 mL of microsome suspension buffer (MSB; 10 mM KH_2_PO_4_/K_2_HPO_4_ buffer, pH 7.8, 0.3 M sucrose) with a teflon-glass homogenizer. Samples were centrifuged again at 231,000 x *g* for 35 min and the pellet was suspended in MSB (2 mL) prior to homogenization using an electric teflon-glass homogenizer.

A two-phase partition medium was produced by adding 1.45 g of polyethylene glycol (3,350) and 1.45 g of dextran to 9.3 mL of MSB and 7.3 mL of NaCl medium (1.17 g of NaCl in 200 mL of MSB). The MSB homogenate was placed in the two-phase partition medium and incubated on ice for 10 min, with periodic shaking. Samples were centrifuged (650 x *g* for 5 min at 4 °C) and the upper phase was transferred to a new partition medium, with this procedure conducted a total of three times. The upper phase was transferred to PM-suspension medium (10 mM MOPS buffer-KOH (pH 7.3), 1 mM EGTA (pH 8.0), 0.3 M sucrose, and 2 mM DTT) and centrifuged at 231,000 x *g* for 35 min at 4 °C. The pellets were then homogenized in PM-suspension medium before repeating the 231,000 x *g* centrifugation. The final pellets were then homogenized on ice in 500 μL of PM-suspension medium using an electric Teflon-glass homogenizer, flash frozen, and kept at -80 °C until use.

Isolations were done four times on independent plant material at each time point.

### Enzyme activity assays

Verification of cell fractionation and PM enrichment was confirmed by enzyme marker assays for the presence of plasma membrane (vanadate stimulated H^+^ ATPase), tonoplast (nitrate stimulated H^+^ ATPase), mitochondria (cytochrome c oxidase), endoplasmic reticulum (NADH cytochrome c reductase), and Golgi apparatus (Triton X-100 stimulated UDPase) in the microsomal and plasma membrane enriched fractions as previously described (Uemura *et al*. 1995). In addition, chlorophyll content was determined in the two fractions to estimate the content of the thylakoid membrane of chloroplasts.

### Tryptic digest and nano-liquid chromatography-mass spectrometry

Samples were electrophoresed on a e-PAGEL HRmini gel (ATTO Products, Tokyo, Japan) to remove non-proteinaceous material and excised for in-gel tryptic digestion as described previously (Takahashi *et al*. 2012; Kamal *et al*. 2020). Tryptic peptides were subsequently desalted and concentrated using a solid-phase extraction C-TIP T-300 (Nikkyo Technos Co., Tokyo, Japan). Peptide solutions were first concentrated with a trap column (L-column Micro 0.3 × 5 mm; CERI, Japan) and then separated with a Magic C18 AQ nano column (0.1 × 150 mm; MICHROM Bioresources, Auburn, CA) using a linear gradient of 5-45% acetonitrile at a flow rate of 500 nL min^-1^. After peptides were ionized by an ADVANCE UHPLC spray source (MICHROM Bioresources, Auburn, CA) at 1.8 KV spray voltage, mass analysis was performed using an LTQ Orbitrap XL mass spectrometer (Thermo Fisher Scientific, Waltham, MA) equipped with Xcalibur software (version 2.0.7, Thermo Fisher Scientific). Full-scan mass spectra were obtained in the range of 400 to 1800 m/z with a resolution of 30,000. Collision-induced fragmentation was applied to the five most intense ions at a threshold above 500. Experiments for each treatment were conducted at least four times.

For semi-quantitative analysis, raw files were analyzed using Progenesis liquid chromatography-mass spectrometry (MS) software (version 4.0, Nonlinear Dynamics, New Castle, U.K.). Peptides were assigned to proteins by MASCOT searching (version 2.3.02, Matrix Science, London, U.K.) against the *Brachypodium* genome (version 2.0). Finally, proteins of interest (those with significantly different abundance profiles as described below) were filtered with analysis of variance (ANOVA, *p* < 0.05) and fold-changes (> 1.5) reported according to normalized peptide intensity.

### Statistical analysis

ANOVA was performed on the raw abundance levels of proteins among the four separate trials of each condition, with *p*-values adjusted via the Benjamini-Hochberg method (Ferreira and Zwinderman 2006) to account for multiple testing. Proteins with an absolute fold-change value > 1.5 (adjusted *p* < 0.05) were classified as differentially abundant. For each protein that differed significantly across treatments, Dunnett’s tests were performed to determine the experimental conditions in which the protein’s abundance differed significantly from the control (NA) condition (*p* < 0.05; |fold-change| > 1.5). Heatmaps were generated for proteins with significantly different abundance from the NA control in at least one treatment, using log_2_-transformed fold-change values. Proteins absent from one or more experimental conditions were not included in heatmaps. All code and scripts used in this study can be found in File S1.

### Protein functional classification and subcellular localization predictions

Descriptions were manually predicted using UniProt (UniProt Consortium), RIKEN Brachypodium FLcDNA database (Mochida *et al*. 2013), BLAST, and through literature searches. PM localizations were predicted using UniProt, TMHMM Server (version 2.0) for transmembrane helices (Krogh *et al*. 2001), GPS-lipid for N-myristoylation/-palmitylation sites, DeepLoc-1.0 (http://www.cbs.dtu.dk/services/DeepLoc-1.0/) (Almagro Armenteros *et al*. 2019), BUSCA (http://busca.biocomp.unibo.it/) (Savojardo *et al*. 2018), WolF PSORT (https://wolfpsort.hgc.jp/) (Horton *et al*. 2007), and known localization of orthologous plant proteins. Localization to other compartments was predicted using Uniprot (UniProt Consortium) and SignalP 5.0 (http://www.cbs.dtu.dk/services/SignalP/) (Almagro Armenteros *et al*. 2019) for localization to the extracellular space, mitochondria, and chloroplasts. Proteins were classified based on the functional categories as described by Bevan *et al*. (1998) and Miki *et al*. (2019).

Comparisons between the *Brachypodium* CA dataset (this study) and a previously reported *Arabidopsis* CA dataset (Miki *et al*. 2019) were conducted by analyzing minimum and maximum fold-changes following two days of CA for both species as well as following six days of CA for *Brachypodium* and seven days of CA for *Arabidopsis*. A pseudo-count of one was added for proteins not detected in NA conditions. *Arabidopsis* proteins were selected based on an ANOVA *p* < 0.05 and max fold-change > 1.5 or min fold-change < 0.67 using data provided in Supplemental Table S1 from Miki *et al*. (2019). Orthologous *Brachypodium* and *Arabidopsis* proteins were identified as reported in the publicly available OMA Browser (Altenhoff *et al*. 2021).

**Table 1.**
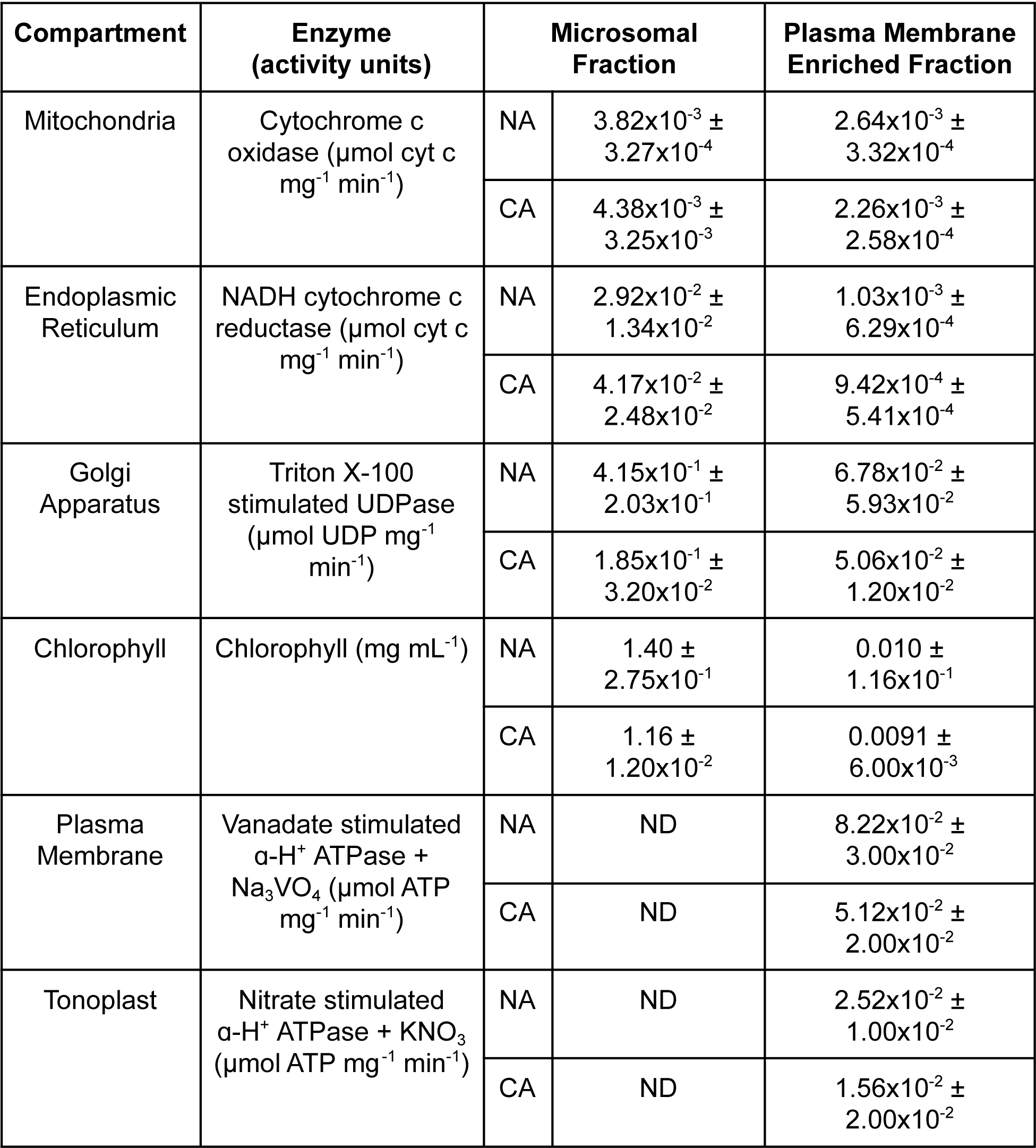
Specific activities of enzymes used as cellular compartment markers for the monitoring of the fractionation of tissue samples collected from *Brachypodium distachyon* either cold-acclimated (CA) for two days at 2 °C, or non-CA (NA), with values not determined (ND) for microsomal fractions of plasma membrane and tonoplasts; experiments were conducted four times with similar results, and as indicated by the standard deviation of the mean shown.

| Compartment | Enzyme<br>(activity units) | Microsomal<br>Fraction |  | Plasma Membrane<br>Enriched Fraction |
| --- | --- | --- | --- | --- |
| Mitochondria | Cytochrome c<br>oxidase ($\mu\text{mol cyt c}$<br>$\text{mg}^{-1} \text{min}^{-1}$) | NA | $3.82 \times 10^{-3} \pm$<br>$3.27 \times 10^{-4}$ | $2.64 \times 10^{-3} \pm$<br>$3.32 \times 10^{-4}$ |
| | | CA | $4.38 \times 10^{-3} \pm$<br>$3.25 \times 10^{-3}$ | $2.26 \times 10^{-3} \pm$<br>$2.58 \times 10^{-4}$ |
| Endoplasmic<br>Reticulum | NADH cytochrome c<br>reductase ($\mu\text{mol cyt c}$<br>$\text{mg}^{-1} \text{min}^{-1}$) | NA | $2.92 \times 10^{-2} \pm$<br>$1.34 \times 10^{-2}$ | $1.03 \times 10^{-3} \pm$<br>$6.29 \times 10^{-4}$ |
| | | CA | $4.17 \times 10^{-2} \pm$<br>$2.48 \times 10^{-2}$ | $9.42 \times 10^{-4} \pm$<br>$5.41 \times 10^{-4}$ |
| Golgi<br>Apparatus | Triton X-100<br>stimulated UDPase<br>($\mu\text{mol UDP mg}^{-1}$<br>$\text{min}^{-1}$) | NA | $4.15 \times 10^{-1} \pm$<br>$2.03 \times 10^{-1}$ | $6.78 \times 10^{-2} \pm$<br>$5.93 \times 10^{-2}$ |
| | | CA | $1.85 \times 10^{-1} \pm$<br>$3.20 \times 10^{-2}$ | $5.06 \times 10^{-2} \pm$<br>$1.20 \times 10^{-2}$ |
| Chlorophyll | Chlorophyll ($\text{mg mL}^{-1}$) | NA | $1.40 \pm$<br>$2.75 \times 10^{-1}$ | $0.010 \pm$<br>$1.16 \times 10^{-1}$ |
| | | CA | $1.16 \pm$<br>$1.20 \times 10^{-2}$ | $0.0091 \pm$<br>$6.00 \times 10^{-3}$ |
| Plasma<br>Membrane | Vanadate stimulated<br>$\alpha\text{-H}^+$ ATPase +<br>$\text{Na}_3\text{VO}_4$ ($\mu\text{mol ATP}$<br>$\text{mg}^{-1} \text{min}^{-1}$) | NA | ND | $8.22 \times 10^{-2} \pm$<br>$3.00 \times 10^{-2}$ |
| | | CA | ND | $5.12 \times 10^{-2} \pm$<br>$2.00 \times 10^{-2}$ |
| Tonoplast | Nitrate stimulated<br>$\alpha\text{-H}^+$ ATPase + $\text{KNO}_3$<br>($\mu\text{mol ATP mg}^{-1} \text{min}^{-1}$) | NA | ND | $2.52 \times 10^{-2} \pm$<br>$1.00 \times 10^{-2}$ |
| | | CA | ND | $1.56 \times 10^{-2} \pm$<br>$2.00 \times 10^{-2}$ |

### Construction of networks

A list of protein accession identifications for all significantly increased and decreased proteins obtained by MS were assembled and used as inputs for STRING (version 11.0) to predict protein-protein interactions (Franceschini *et al*. 2016; Szklarczyk *et al*. 2019) for CA2 and CA6 timepoints. A predicted network was prepared and exported to Cytoscape (version 3.8.1) for further modification.

Additional protein metadata was input into Cytoscape including corresponding log_2_ fold-change values which were assigned to node fill mapping.

To construct a stress response meta-analysis network, individual protein accession identifications were subjected to literature searches (performed to 1/1/2021) and annotated according to their protein descriptions and involvement in stress response pathways (File S2). Proteins with no reported involvement in stress responses were omitted. The dataset was then input into Cytoscape with and log_2_ fold-changes were again selected as node fill mapping as described previously. All networks were centred in the plot area and exported as Scalable Vector Graphics (SVG) files where further modification was performed and legends added in Inkscape (version 0.92.2). Interactive versions of each network were additionally exported as full webpages for viewing in any modern web browser as HTML files with all metadata.

## RESULTS

### Freezing tolerance and accumulation of sugars

Electrolyte leakage assays were conducted to assess leaf membrane integrity of CA and NA *Brachypodium* following exposure to temperatures ranging from -2 °C to -10 °C (Figure 1A). Electrolyte leakage was significantly reduced compared to NA plants following two or more days of CA at temperatures between -4 and -10 °C. Increasing CA duration up to 28 days did not further reduce electrolyte leakage (Figure S1). These results are consistent with earlier reports demonstrating that *Brachypodium* achieved peak freezing tolerance after two days of CA at 2 °C (Colton-Gagnon *et al*. 2014; Bredow *et al*. 2016).

**Figure 1.**
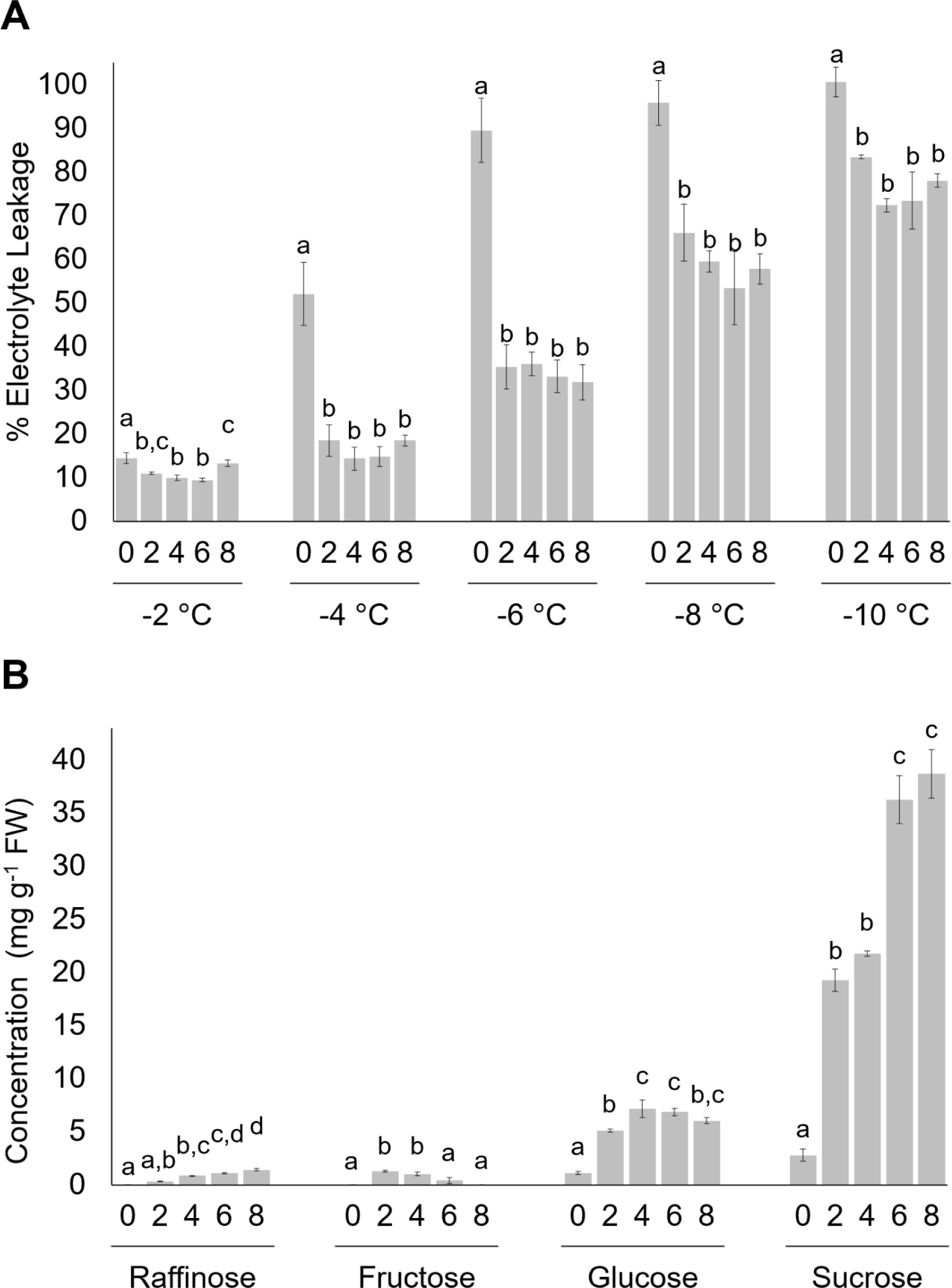
Impact of duration of cold acclimation on *Brachypodium distachyon* freezing tolerance. **(A)** Electrolyte leakage assays conducted using leaf tissue from non-acclimated and cold-acclimated wildtype ecotype *Bd*21. Days of cold acclimation and temperatures are shown. Samples were subjected to the low-temperatures indicated at the rate of 1 °C every 15 min before assaying for electrolyte leakage (%). **(B)** Accumulation of soluble sugars in leaf tissue from non-acclimated and cold-acclimated wildtype ecotype *Bd*21. Days of cold acclimation and soluble sugar type is shown in four clusters of histograms and expressed as concentration in fresh weight (FW) of leaves. Four biological replicates were conducted for all assays (n = 10) and ANOVA and post-hoc Tukey tests were performed. Error bars represent standard error of the mean. Letters above the histograms indicate statistically significant groups (*p* < 0.05) with separate analyses for each temperature and soluble sugar.

The accumulation of osmoprotectant sugars, or compatible solutes, mitigates the effects of freeze-induced cellular dehydration and is associated with increased freezing tolerance (Tarkowski and Van den Ende 2015). Here, the accumulation of sucrose increased from 2.8 mg g^-1^ in NA leaf tissue to 19.3 mg g^-1^ and 38.7 mg g^-1^ fresh weight (FW) in CA2 and CA8, respectively (Figure 1B; Figure S2). Glucose increased 4.6-fold to 5.1 mg g^-1^ in CA2 leaves and remained relatively stable for the duration of cold treatment. Raffinose showed a similar profile to sucrose with consistent increases in abundance throughout CA, reaching a maximum of 1.3 mg g^-1^ after eight days. In contrast, fructose accumulation peaked in CA2 leaves (1.3 mg g^-1^ FW), and steadily declined in the remainder of the cold treatment groups. Thus, CA in *Brachypodium* is characterized by the early accumulation of glucose, raffinose, fructose and sucrose, coincident with significantly reduced electrolyte leakage.

Under longer acclimation regimes most soluble sugars increased in abundance, with the exception of fructose, and did not correspond to further changes in membrane integrity.

### Cold acclimation treatment time and quantifying recovered proteins

PM-associated proteins were analyzed in NA *Brachypodium* and following CA (CA2, CA4, CA6 and CA8). PM fractions were prepared by a method adapted for *Arabidopsis* (Kamal *et al*. 2020) with the effectiveness of fractionation and enrichment of PMs in *Brachypodium* validated using marker enzymes in NA and CA2 samples (Table 1). PM fractions of both NA and CA samples showed an overall decrease in mitochondria, endoplasmic reticulum, Golgi, and chloroplasts relative to the microsomal fractions. In contrast, vanadate-stimulated H^+^ ATPase activity, associated with the PM, showed an overall enrichment in PM fractions, with a small amount of contamination from the tonoplast, indicated by nitrogen-stimulated H^+^ ATPase activity (Table 1).

PM-enriched fractions were subjected to nano-liquid chromatography-MS for the label-free quantification of proteins, resulting in the identification of a total of 1,349 unique peptides corresponding to known proteins (File S3). Many of these (848 or 63% of the total) showed a significant change in relative abundance (*p* < 0.05, |fold-change| > 1.5) during cold treatment compared to NA plants. Of these proteins, 334 (39%) significantly increased > 1.5-fold in relative abundance (File S4) at one or more time points, with more (522; 62%) showing significant decreases > 1.5-fold (File S5). In line with the cold-induced membrane protection observed in electrolyte leakage assays (Figure 1), several protein families associated with cold stress were identified in our MS analysis including sugar transporters (sucrose transporter SUT1-like protein; Bradi1g73170.1; Tarkowski and Van den Ende 2015), dehydrins (dehydrin COR410-like; Bradi3g51200.1; Liu *et al*. 2017), and ROS scavengers (catalase; Bradi1g76330.1; Yousefi *et al*. 2018), and served to increase confidence in the analysis.

To better visualize changes in the proteome following CA, a heat map was generated using log_2_ transformed fold-changes at each treatment point for all proteins with significant changes in protein abundance (Figure 2A). Of the 334 proteins that showed relative increases in abundance after CA compared to NA plants, 57 (17%), 129 (39%), 224 (67%), and 205 (61%) were increased in CA2, CA4, CA6 and CA8 plants, respectively. The majority showed moderate increases over > 1.5-fold compared to control levels, with some notable exceptions particularly after longer cold treatments. For example, after two days of CA, 7% and 3% of the proteins were one and two orders of magnitude more abundant, respectively, than in NA samples, but with higher levels achieved by more proteins at CA6. Overall, fold-changes for the 848 tracked proteins showed that the CA6 and CA8 treatment profiles were more alike than the similar CA2 and CA4 sample profiles. Using the CA2 proteome and the CA6 proteome as typifying the early and a sustained response, respectively, showed distinctive frequency distributions in the numbers of proteins that were relatively more or less abundant than proteins in the NA plants (Figure 2B,C). Although as indicated, the relative abundance of more proteins were affected by the longer low-temperature treatment, in both CA2 and CA6 treatment-groups there were about twice as many proteins that decreased *vs.* increased in relative abundance (*i.e.* there were 52 and 97 proteins that increased and decreased in abundance, respectively in CA2, with corresponding numbers of 224 and 456 for CA6; Figure S3).

**Figure 2.**
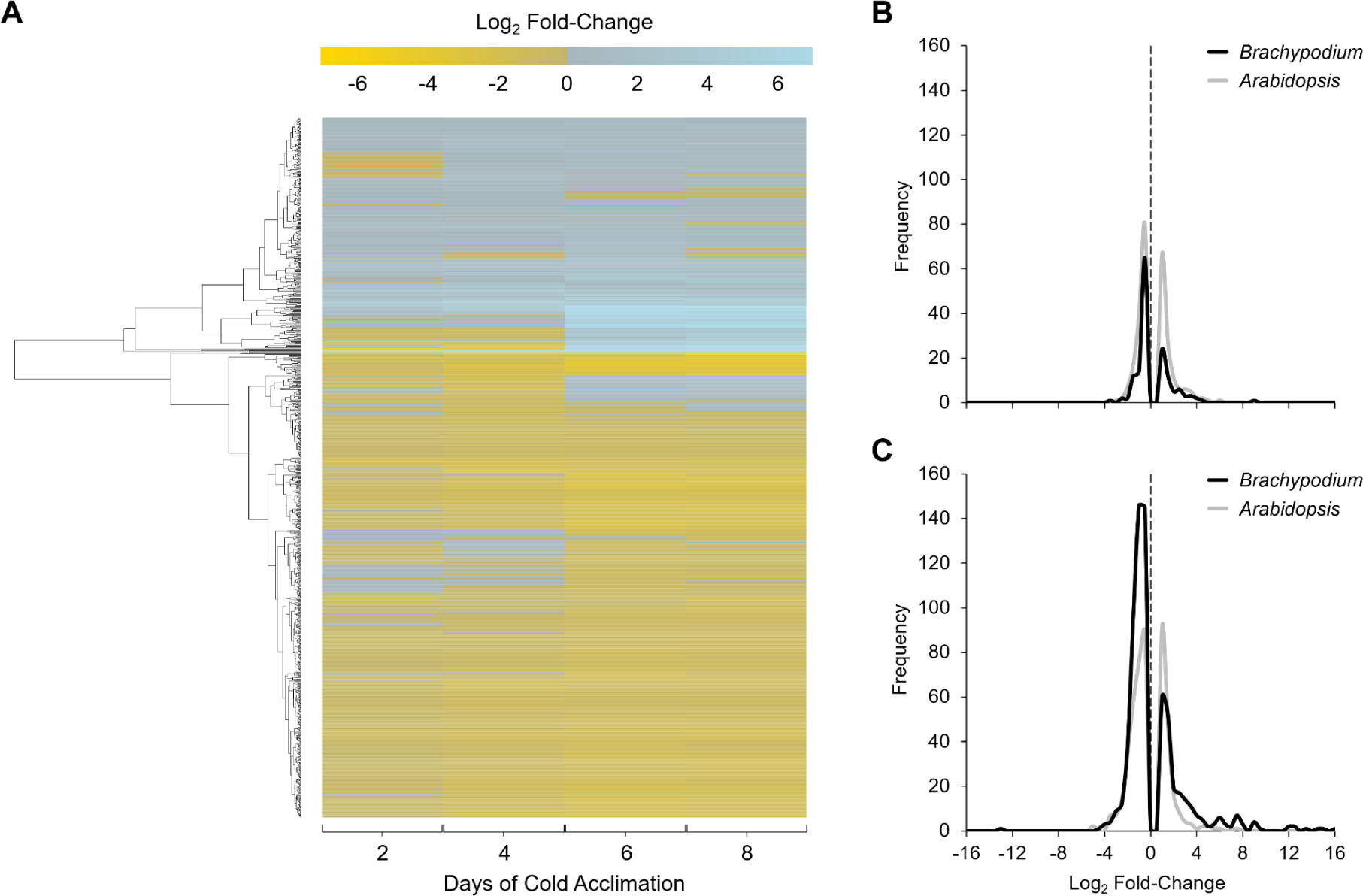
Global impact of cold acclimation on the *Brachypodium distachyon* PM proteome. **(A)** A heatmap showing the 848 proteins in the PM enriched fractions with significant changes in relative abundance between non-acclimated and cold-acclimated samples following two, four, six, or eight days at 2 °C. Data are mean fold-changes relative to non-acclimated plants based on four replicates, and are presented as log_2_ transformed values. Hierarchical clustering analysis was used to group proteins displaying similar abundance proteins, and the results are presented by the dendrogram along the left-hand side of the figure. **(B,C)** Frequency distribution of log_2_ fold-changes shown for proteins with significant changes in relative abundance at **(B)** early cold acclimation (two days for *B. distachyon* and two days for *Arabidopsis thaliana*) and **(C)** sustained cold acclimation (six days for *B. distachyon* and seven days for *A. thaliana*). Log_2_ fold-change values of 0 are highlighted with a dotted line. Data was plotted using fold-change value bin widths of 0.5. Raw *A. thaliana* data was obtained from Miki *et al*. (2019) and re-analyzed to obtain fold-changes matching thresholds used in this manuscript of *p* < 0.05 and a fold-change > 1.5 or < 0.67 as opposed to > 2.0 and < 0.5 employed by Miki *et al*. (2019).

Proteins were functionally annotated and then categorized using UniProt (UniProt Consortium 2019), BLAST, and through the literature (Figures 3, 4, S6-S9). Many proteins that increased in relative abundance during CA were related to energy and metabolism (16 in CA2 and 55 in CA6 plants) such as PM ATPase (Bradi5g24690.1; Nguyen *et al*. 2018). They also represented disease and defense functions (14 in CA2 and 29 in CA6 plants) exemplified by dehydrin COR410-like (Bradi3g51200.1; Chiappetta *et al*. 2015), and transport and signal transduction affiliates (nine in CA2 and 25 in CA6 plants) such as heavy metal-associated isoprenylated plant protein 27 (Bradi1g70700.1; de Abreu-Neto *et al*. 2013). Proteins that increased in abundance with the greatest fold changes are listed in Figure 3.

**Figure 3.**
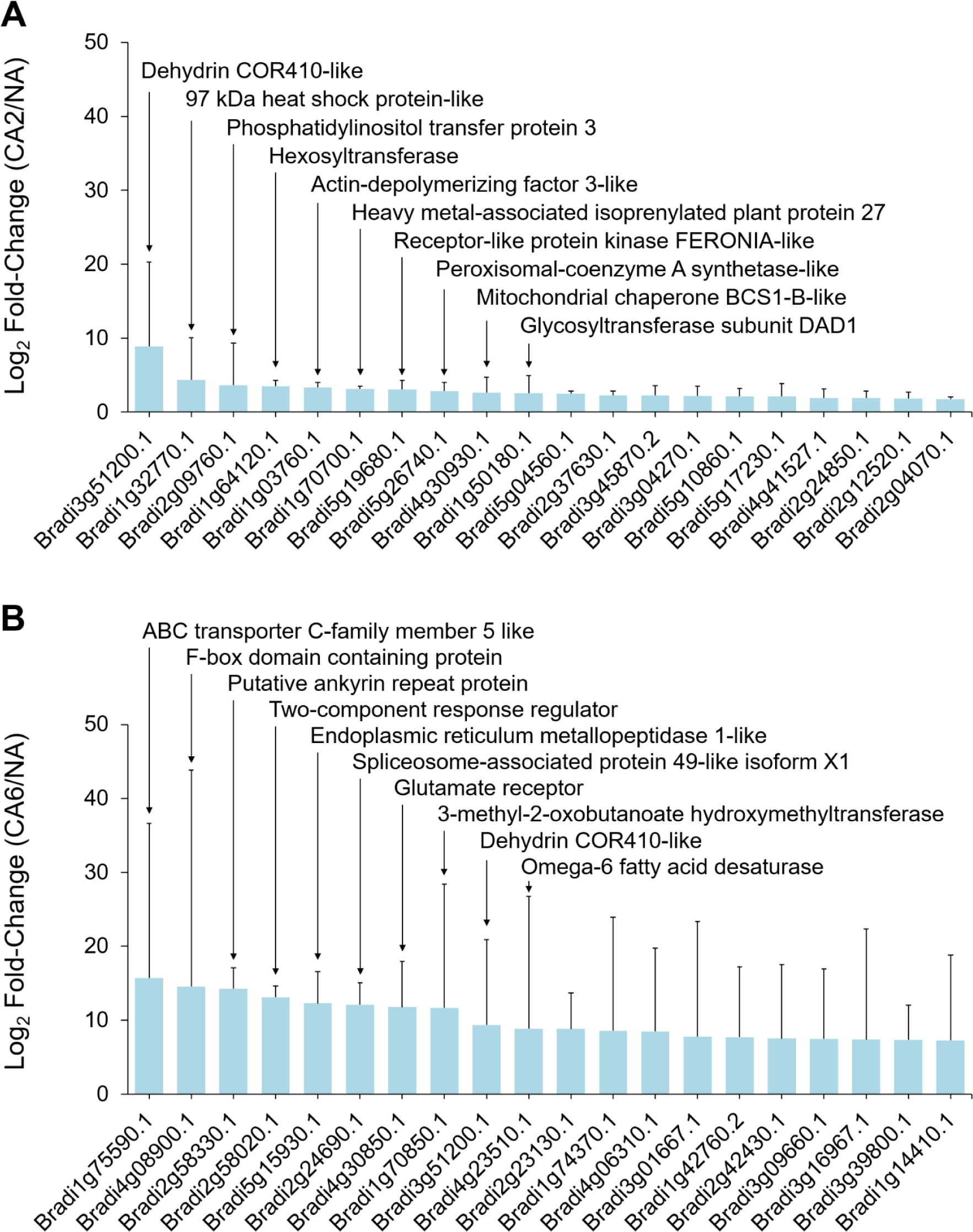
*Brachypodium distachyon* proteins undergoing the greatest fold increases in relative abundance during cold acclimation. Data is shown for **(A)** two days and **(B)** six days of cold acclimation. Values are the average of four replicate trials and protein annotations were predicted as previously described. Error bars represent standard error of the mean. A pseudo-count of one was added to the non-cold-acclimated value for proteins that were not detected under that experimental condition. The ten proteins with the greatest fold relative increases are labelled accordingly.

Despite our division of an early (CA2 and CA4 plants) and a sustained response to low-temperature treatment (CA6 and CA8 plants), there were similarities in the cellular functions in both groups.

The total of 522 proteins that decreased in relative abundance after CA when compared to NA plants showed a frequency profile that increased over time: 97 (19%), 332 (64%), 456 (87%), and 421 (81%) proteins showed significant relative decreases at CA2, CA4, CA6 and CA8, respectively (Figure 2, Figure S3). For the most part (61%), the identified proteins showed relative decreases in abundance that were at, or just modestly greater than, 1.5-fold, but approximately 1% and <1% showed decreases that were one and two orders of magnitude greater, respectively. Proteins that decreased in relative abundance with the greatest fold changes are listed (Figure S4), and according to functional groupings (Figure 4). The most notable decreases in relative abundance were in proteins with likely functions in energy metabolism (24 in CA2 and 97 in CA6 plants), including cytochrome b_5_, Bradi2g62010.1 (Gou *et al*. 2019) and probable L-gulonolactone oxidase 4, Bradi3g13700.1 (Maruta *et al*. 2010). Proteins belonging to other functional classifications also decreased in relative abundance after low-temperature treatment. These included those associated with disease and defense (13 in CA2 and 68 in CA6 plants) as well signal transduction (seven in CA2 and 51 in CA6 plants). Not all of the identified proteins are known to localize to the PM, as discussed below.

**Figure 4.**
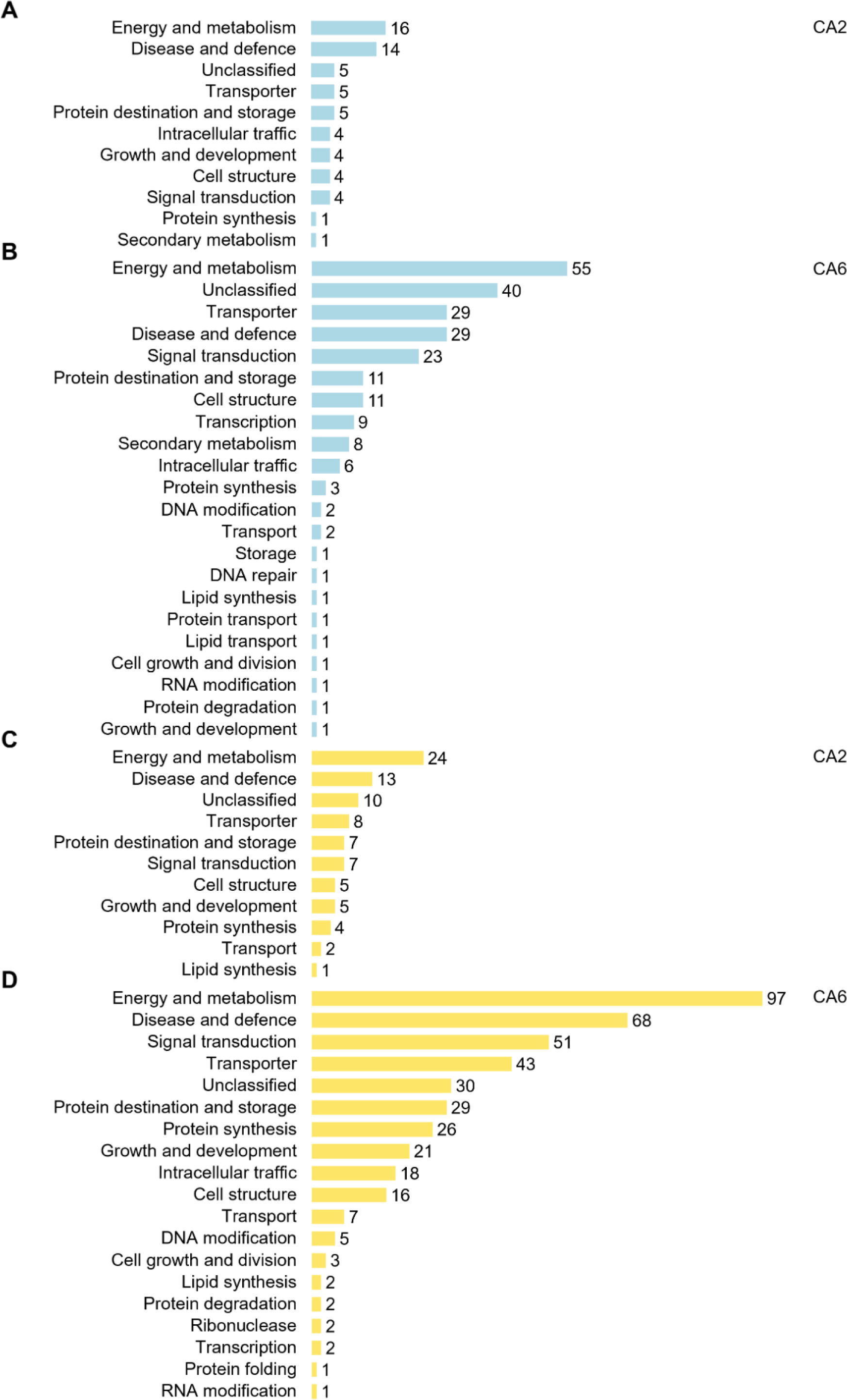
Functional analysis of proteins whose relative abundance was impacted by two days and six days of cold acclimation. Predicted functional categories of proteins that **(A)** increased in relative abundance at two days and **(B)** six days and **(C)** decreased in relative abundance at two days and **(D)** six days (|mean fold-change| > 1.5, *p*-value ≤ 0.05, based on four replicates) compared to non-acclimated controls. Functional descriptions were manually predicted using UniProt, RIKEN, *Brachypodium* FL cDNA database, BLAST, and literature searches. Scales are all equal.

### Identification of proteins involved in the two phase cold-induced response

As indicated, the numbers of significant changes in PM-associated protein relative abundance were highest at CA6, although electrolyte leakage assays indicated that CA2 plants were similarly protected from freezing damage (Figure 1A; Figure S1). This suggests that early CA-mediated PM changes, including changes in the proteome, are sufficient for membrane protection. As indicated, the impact on the proteome could be grouped as an early (CA2 and CA4) and a sustained (CA6 and CA8) response and thus a categorical analysis of the annotated proteome focused on proteins increased in abundance in CA2 and CA6 plants to represent these two phases (Figure 2 and Figure 4). In CA2 plants, the protein with the greatest relative increase (483-fold) was a predicted dehydrin COR410-like protein (Bradi3g51200.1), which increased an order of magnitude more than the second and third most abundantly increased proteins, a 97 kDa heat shock protein-like, Bradi1g32770.1 and a phosphatidylinositol transfer protein, Bradi2g09760.1, respectively (Figure 3A; File S8). All of these proteins likely play important roles in ensuring membrane integrity. The accumulation of the COR410-like protein remained elevated with respect to NA plants in CA6 plants, however, eight other proteins showed greater relative fold-increases at the later time (Figure 3B). Thus, the early proteome response to low-temperatures, coincident with CA-induced membrane protection, was followed by a later response that included a large number of additional proteins.

The three proteins with the greatest relative fold-increases in CA6 plants were an ABC transporter C-family 5 like protein (Bradi1g75590.1), an F-box domain containing protein (Bradi4g08900.1), and a putative ankyrin repeat protein (Bradi2g58330.1) that showed a >19,000-fold increase in relative abundance over the level in NA plants (Figure 3B). Other proteins with large increases in relative abundance (> 1.5-fold) following six days of CA included ROS scavengers (catalase, Bradi1g76330.1; phospholipid hydroperoxide glutathione peroxidase 1, Bradi1g47140.1), heat shock proteins or HSPs (97 kDa heat shock protein-like, Bradi1g32770.1; heat shock cognate 70 kDa protein-like, Bradi1g66590.1), sugar transporters (bidirectional sugar transporter SWEET1b-like, Bradi2g24850.1; sucrose transport protein SUT1-like, Bradi1g73170.1), dehydrins (the mentioned dehydrin COR410-like, Bradi3g51200.1; dehydrin COR410-like, Bradi5g10860.1), heavy metal-associated isoprenylated plant proteins or HIPPs (heavy metal-associated isoprenylated plant protein 27, Bradi1g70700.1; heavy metal-associated isoprenylated plant protein 20, Bradi3g04270.1), phospholipase D (Bradi4g36800.1), and a glutamate receptor (Bradi4g30850.1). As a group these could function to support long-term survival under conditions of stress. In addition, there were five proteins that were only identified following CA, which are predicted to have roles in transcription (RRM domain-containing protein, Bradi2g24690.1; Lorković and Barta 2002), signal transduction (two-component response regulator, Bradi2g58020.1; Lohrmann and Harter 2002), defense signaling (putative ankyrin repeat protein, Bradi2g58330.1; Yang *et al*. 2012), and cold perception (glutamate receptor, Bradi4g30850.1; Gong *et al*. 2019), as well as an unclassified protein (ER metallopeptidase 1-like, Bradi5g15930.1; Marino and Funk 2012). The complete dataset of annotated CA6 increased proteins is available in File S9.

Concomitant with the increased abundance of many PM proteins, those that decreased in relative abundance by two days of CA included enzymes likely required for nutrient uptake, PM H^+^ ATPases (PM ATPase, Bradi5g24690.1 and PM ATPase 1-like, Bradi1g54847.1; Morsomme and Boutry 2000). By CA6 some of these further decreased in relative abundance and were joined by additional PM H^+^ ATPases, consistent with reports from cold-stressed *Arabidopsis* (Muzi *et al*. 2016). Likewise, other PM ATPases decreased in relative abundance after two days (V-type ATPase, Bradi1g67960.1 and AAA+ ATPase, Bradi1g48010.1), and more after six days of CA (another AAA+ ATPase, Bradi1g36830.1; V-type proton ATPase subunit F, Bradi2g39610.1; and Obg-like ATPase, Bradi3g17680.1), further suggesting an early and sustained response to low-temperature exposure at least with regard to the numbers of impacted proteins. It is important to note that although many of the

PM-associated proteins that decreased in relative abundance have been categorized to a single function, most notably in energy and metabolism (Figure 4), some may have other documented roles.

Protein localization algorithms (DeepLoc-1.0, BUSCA, WolF PSORT) predicted that 46% (26/57) of proteins at CA2 and 39% (88/224) of proteins at CA6 that increased in relative abundance were associated with the PM, which is not surprising given the technique designed to enrich for PM-associated proteins and highlighting the difficulty in the recovery of hydrophobic PM proteins. In contrast, 42% (41/97) of the proteins at CA2 and 38% (175/456) of those identified at CA6 that showed decreases in relative abundance were predicted to localize to the cytosol (Files S6 and S7). Although highly abundant cytosolic proteins could have contaminated our PM preparations, some of these proteins may in fact loosely associate with the PM during normal growth and weaken as the PM is restructured in response to low-temperature. Links to the PM are known to vary with reversible lipid interactions (Marmagne *et al*. 2004; Komatsu *et al*. 2006) although this remains to be experimentally determined for proteins identified in our analysis.

### Crosstalk between stress response networks

Low-temperatures can result in an increasing vulnerability to effects of additional biotic and abiotic stresses in plants (Fujita *et al*. 2006; Rejeb *et al*. 2014), and thus CA regimes may result in abundance changes in seemingly unrelated stress response proteins to safeguard against the effects of combinatorial stresses. In order to gain a better understanding of crosstalk between stress networks at the PM, a network map was constructed using stress-related proteins that increased or decreased in relative abundance compared to NA plants after two days of CA (Figure 5; see File S10 for an interactive version of the network, and File S2 for the underlying data). As expected, the cold stress node showed the largest number of interactions with other stress nodes at 41 proteins, including 20 interactions that were also associated with drought, five with osmotic stress, and 15 with salt stresses. These proteins included those with predicted functional categories of energy and metabolism (12), disease and defense (11), transporters (six), and signal transduction (four). A secondary cluster, the pathogen stress node with 39 proteins, showed interactions with other stress nodes, many which overlapped with those seen with cold stress. Due to the significantly higher number of proteins, it was not practical to construct a corresponding network map for CA6 plants, but functional analysis of the proteins suggests that interactions between abiotic stresses would be retained.

**Figure 5.**
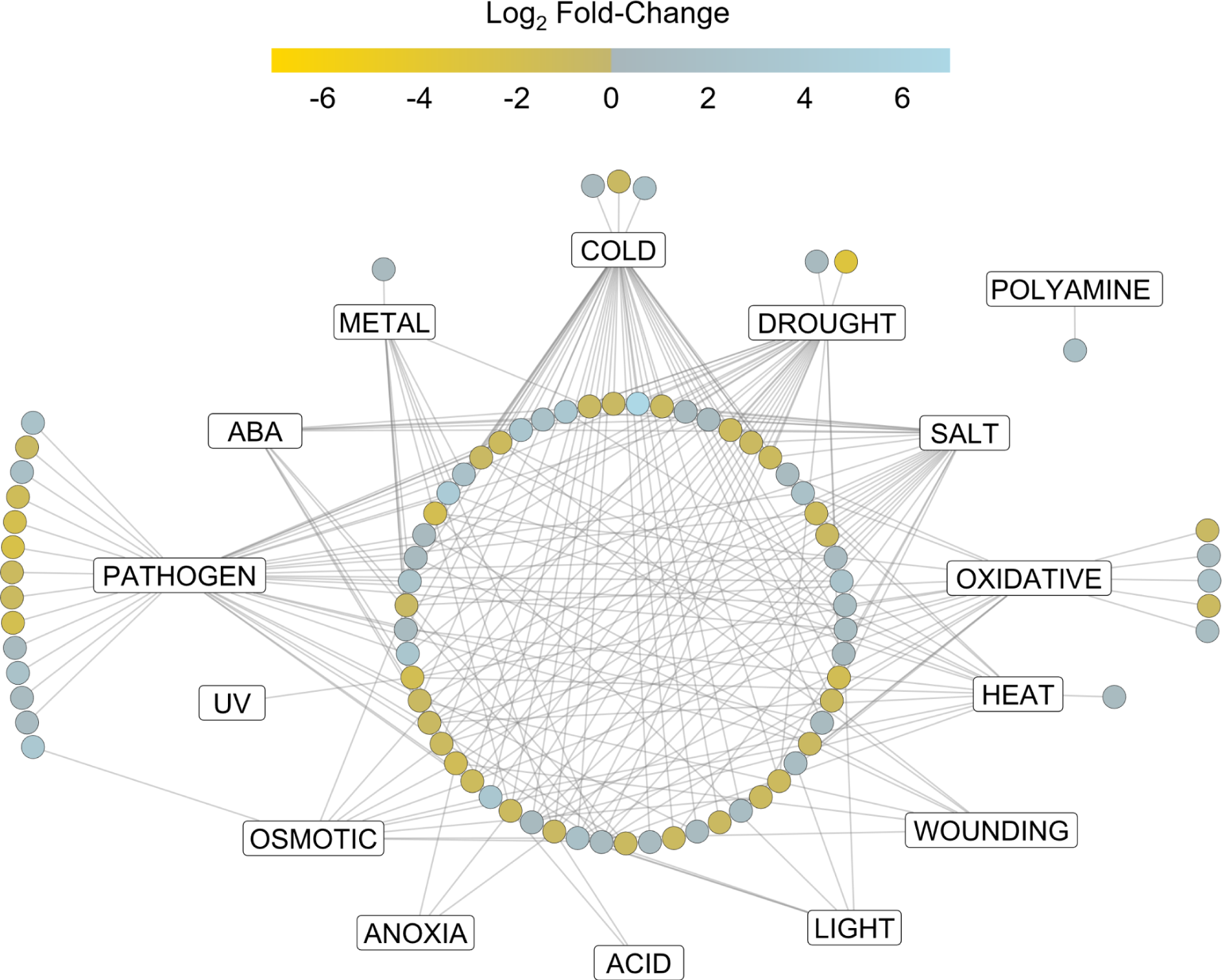
Network illustrating involved stress pathways and crosstalk between proteins significantly changed in relative abundance at two days of cold acclimation. Rectangle nodes represent functional protein stress response pathways while circular nodes represent individual proteins. Edges connecting protein nodes to stress response nodes represent validated functional associations from the literature. Protein function in the stress pathways were validated via Uniprot and the literature, and networks were built using Cytoscape version 3.8.1. Node colour represents log_2_ fold-change of the protein in the cold-acclimated (two days) versus non-acclimated plasma membrane proteomes. An interactive version of the network with encoded metadata is available in File S10. Click on a node to see individual protein accession ID, protein description, log_2_ fold-change, predicted functional categories, and degrees of interaction. Click on an individual edge to see the literature for the specific protein-stress response relationship.

In order to better understand cellular physiology under cold stress conditions, protein-protein interaction networks were predicted *in silico* using STRING (Franceschini *et al*. 2016; Szklarczyk *et al*. 2019). Again, the proteome of CA2 and CA6 plants were chosen to represent early and sustained responses to low-temperature. In CA2 plants, proteins largely associated with protein synthesis, with energy and metabolism and disease and defense, interacted together and with each other in interaction clusters, suggesting that low-temperature exposure initiates metabolic changes, which would not be unexpected (Figure 6; see File S11 for an interactive version of the network with metadata). For example, a predicted subunit theta of T-complex protein 1, an ATP-dependent molecular chaperone protein that assists in protein folding (Bradi4g15450), showed 20 interactions with other proteins and these were primarily heat shock proteins, ribosomal subunits, and actins. Similar clusters were retained in the protein interaction network of CA6 plants, but involved many more protein-protein interactions, consistent with the increased number of differently abundant proteins at this later time (Figure 7 and File S12).

**Figure 6.**
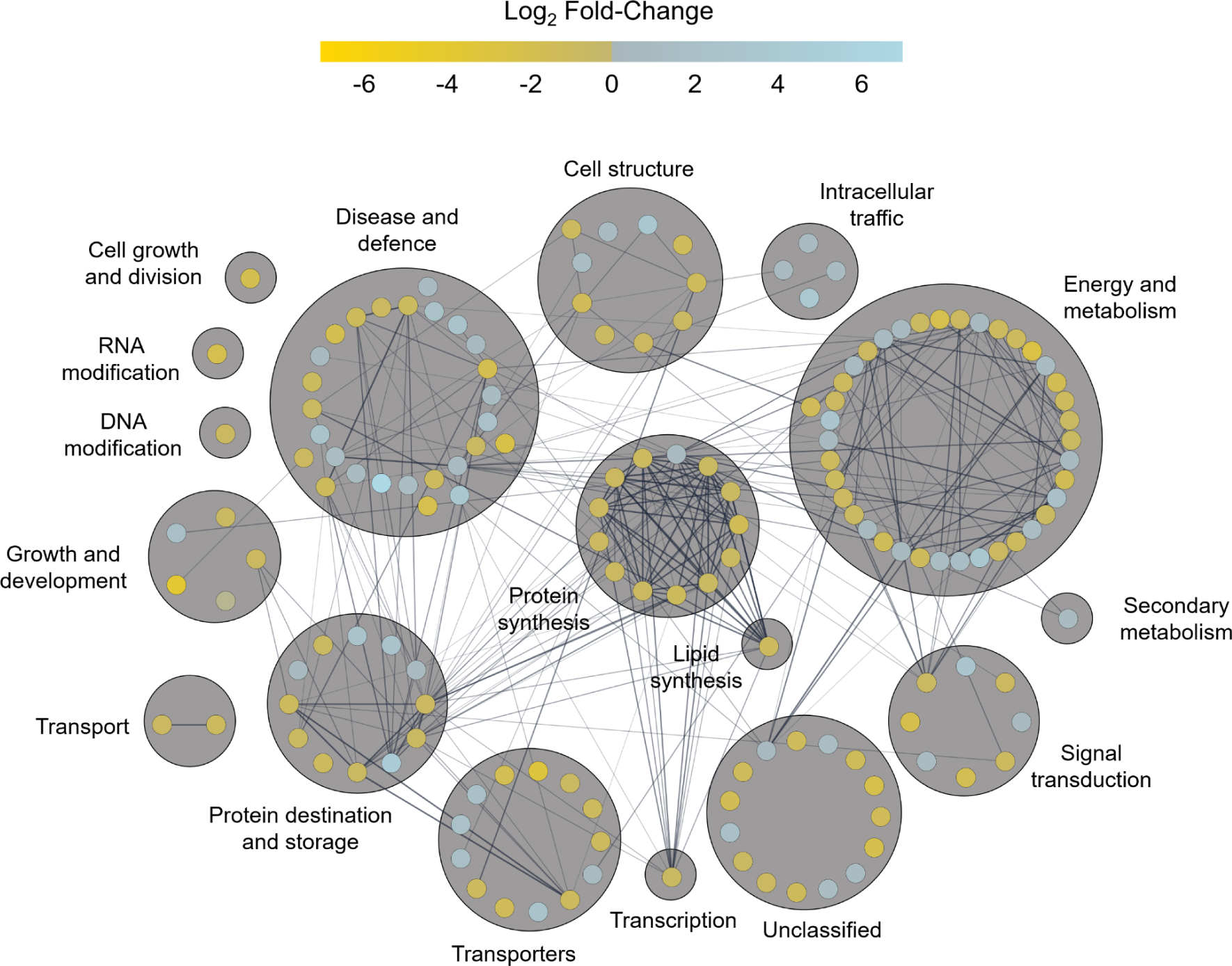
Predicted protein-protein interaction network of all proteins at two days of cold acclimation. Nodes represent individual proteins (154) with node colour representing the respective log_2_ fold-change of the protein in the cold-acclimated (two days) versus non-acclimated plasma membrane proteomes. Edges between connected nodes represent predicted protein-protein interactions. Nodes are grouped based on predicted functional categories and labelled accordingly. Nodes on the outer circle of a main functional group have predicted functions in secondary categories. Interactions were predicted via STRING version 11 and the network was prepared using Cytoscape version 3.8.1. Darker and thicker edges between nodes represents stronger supporting data for the respective interaction. An interactive version of the network with encoded metadata is available in File S11. Click on a node to see individual protein accession ID, protein description, log_2_ fold-change, functional categories, and degrees of interaction.

**Figure 7.**
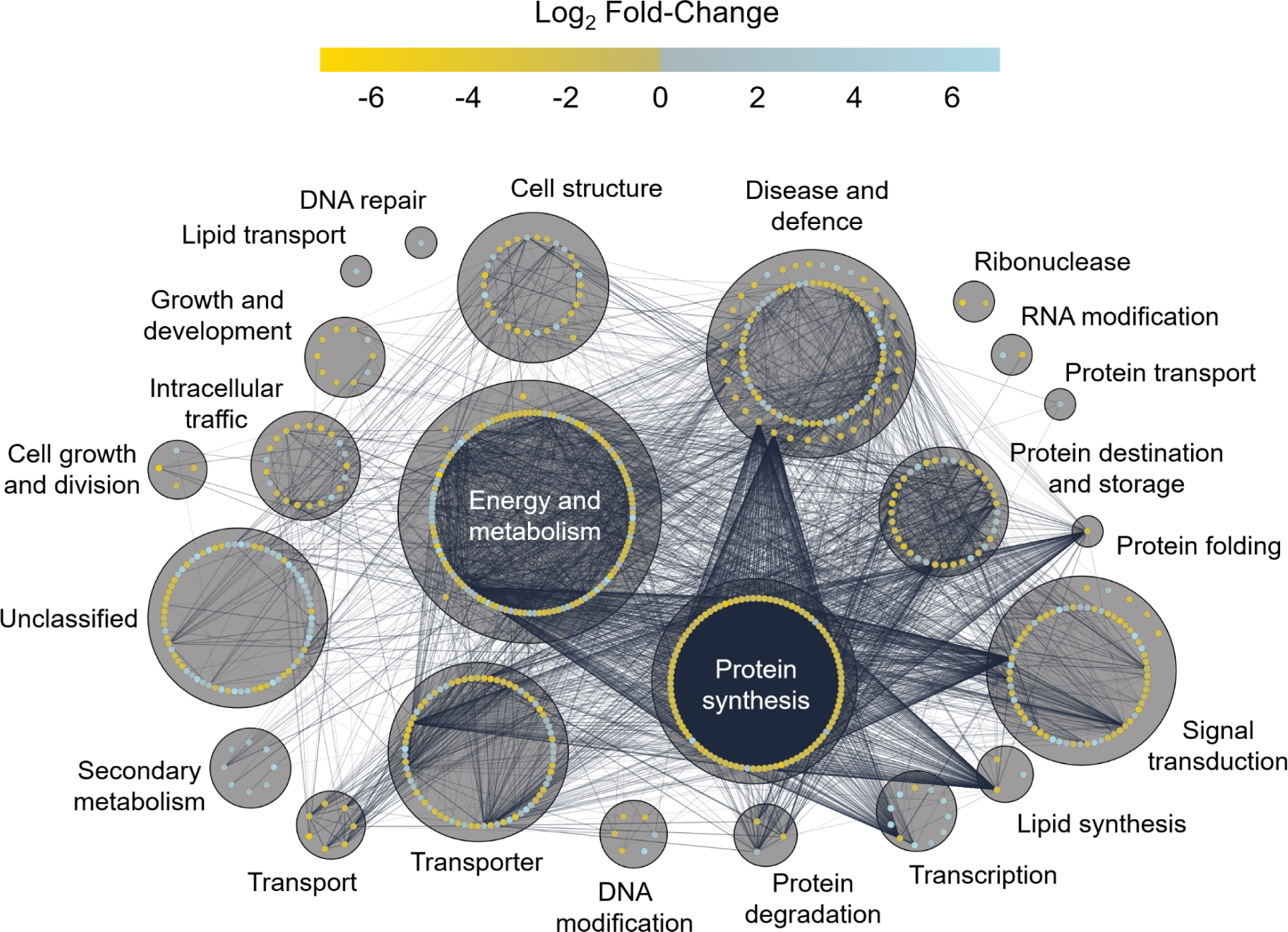
Predicted protein-protein interaction network of all proteins at six days of cold acclimation. Nodes represent individual proteins (680) with fill colour representing the respective log_2_ fold-change of the protein in the cold-acclimated (six days) versus non-acclimated plasma membrane proteomes. Edges between connected nodes represent predicted protein-protein interactions between connected proteins. Nodes are grouped based on predicted functional categories and labelled accordingly. Nodes on the outer circle of a main functional group have predicted functions in secondary categories. Interactions were predicted via STRING version 11 and the network was prepared using Cytoscape version 3.8.1. Darker and thicker edges between nodes represents stronger supporting data for the respective interaction. An interactive version of the network with encoded metadata is available in File S12. Click on a node to see individual protein accession ID, protein description, log_2_ fold-change, functional categories, and degrees of interaction.

STRING analysis predicted interactions between Suppressor of G-Two allele of SKP1 (SGT1, Bradi2g44030.1) with HSP90s (Bradi1g30130.1; Bradi1g32770.1) and HSP70s (Bradi1g66590.1; Bradi2g30660.1) at CA2 (File S11). Interactions between SGT1, HSP90, and HSP70 regulate microbial disease resistance in a number of plant species including *Arabidopsis* (Takahashi *et al*. 2003; Azevedo *et al*. 2006; Noël *et al*. 2007; Spiechowicz *et al*. 2007) and barley (Azevedo et al. 2002; Shen *et al*. 2003; Hein *et al*. 2005; Chapman *et al*. 2021), suggesting that immune signaling pathways are upregulated in response to low-temperature treatment. In CA6 plants, such predicted interactions were conserved but also expanded (File S12). These involved additional HSPs and other immune-related proteins, including stromal HSP70 (Bradi2g30560.1), HSP81-1s (Bradi3g39590; Bradi3g39620.1), activator of HSP90 (Bradi3g37790.1), respiratory burst oxidase homolog protein B-like (Bradi2g12790.2), HSP70-HSP90 organizing protein (Bradi3g50110.1), and two peptidyl-prolyl cis-trans isomerases (Bradi1g34750.1; Bradi2g39950.1) with known functions in immune response (Mokryakova *et al*. 2014). Interestingly, the respiratory burst oxidase homolog protein B-like, which is involved in pathogen resistance in *Arabidopsis* (Hawamda *et al*. 2020), shows additional predicted interactions with calcium-dependent protein kinase 13 (Bradi5g19430.1), mitogen-activated protein kinase (Bradi1g65810.1), catalase (Bradi1g76330.1), and an ammonium transporter (Bradi2g22750.1), further suggesting the activation of immune signaling pathways as a result of cold stress.

A single example of a protein with many interactions at CA6, allene oxide synthase 3 (Bradi3g08250.1), involved in the biosynthesis of jasmonic acid and plant defense (Farmer and Goossens 2019), showed 137 interactions with other proteins associated with functions in protein synthesis, as well as disease and defense. The correlation of time at low-temperature and the number of protein interactions was also reflected in the 6.4-fold increase in the mean number of interactions shown by each protein depicted in the CA2 (3.3) and CA6 (21) networks. Overall, a short period of CA resulted in an interaction network containing 154 nodes as individual proteins, and 252 edges, representing predicted interactions. This increased to 680 nodes and 7,152 edges in CA6 plants (Figures 6 and 7; Files S11 and S12). The increasing complexity of protein-protein interactions and the increased number of cross-talk pathways as low-temperature exposure continues suggests that after the establishment of the early response, the cold resistance phenotype is supported by the proteins associated with the sustained response.

## DISCUSSION

Our motivation to understand PM proteome changes associated with CA in the model monocot *Brachypodium* was the need to address temperature-related challenges to food security linked to climate change. Our results suggest that freeze-tolerance in this species is a dynamic process, with an early frost-resistance response that can be achieved within two days of CA, but thereafter, additional changes to the PM occur four or more days later that presumably allows for a sustained response for low-temperature survival. The presence of these two phases of freezing tolerance is supported by multiple lines of evidence, most notably: i) the acquisition of PM protection in CA2 plants, ii) the rapid accumulation of sucrose by CA2, followed by further sucrose accumulation after six days, and iii) changes in relative protein abundance, demonstrated by heatmaps, functional profiles, as well as network analysis generated from the MS protein discovery that could be divided into two groups: CA2-CA4 and CA6-CA8 (Figures 1-7).

### The early response to cold acclimation

An early or rapid response to low-temperature is sufficient to maintain PM integrity as assessed by electrolyte leakage. Of the 1,349 PM-proteins identified by nano-liquid chromatography-MS, the relative abundance of 57 PM proteins increased > 1.5-fold after two days of CA, and are likely candidates for supportive roles in ensuring membrane integrity at low-temperatures. Consistent with this hypothesis, the vast majority (88%) of these proteins also increased in relative abundance in initial *Brachypodium* experiments that examined the NA and CA2 PM proteome using an acclimation temperature of 4 °C (File S13) demonstrating a conserved and repeatable proteomic response, coincident with membrane integrity. Functional annotation and categorization of the proteins suggested that the early response includes a global shift in proteins from those associated with energy and metabolism, as well as growth and development, identified in the NA PM proteome towards the accumulation of stress-related proteins after CA. Indeed, proteins identified in our two experimental CA *Brachypodium* PM proteomes were fairly consistent with those identified for the CA *Arabidopsis* PM proteome (Miki *et al*. 2019). For example, transporters and signal transduction proteins, including kinases, increased in relative abundance coincident with PM freeze protection in *Arabidopsis* following two days of CA. For *Brachypodium* proteins that significantly decreased and increased in relative abundance (27% and 13%, respectively), at least one ortholog was found to change in abundance in the same direction in the CA *Arabidopsis* PM proteome (Miki *et al*. 2019). When the reverse analysis was performed, 37% and 15% of the significantly decreased and increased proteins, respectively, from the CA *Arabidopsis* PM proteome had an ortholog in our corresponding dataset that changed in abundance in the same direction. This result suggests at least a partially conserved response to low-temperature between *Brachypodium* and *Arabidopsis* at the PM. It is not surprising that they are not identical considering their evolutionary distance and the differences in freezing tolerance as determined by their capacity to survive low-temperatures after CA, with *Arabidopsis* at -6 to -11 °C, depending on the accession, *vs. Brachypodium* at -12 °C; (Kaplan *et al*. 2004; Hannah *et al*. 2006; Mayer *et al*. 2020).

Proteins with the highest increases in relative abundance are obvious candidates to assist in the protection of vulnerable PMs. A striking 483-fold elevation in relative abundance at CA2 was observed for COR410-like (Bradi3g51200.1), a dehydrin family cold-regulated protein. Transcripts encoding this protein have previously been shown to accumulate after low-temperature exposure in *Brachypodium* (Mayer *et al*. 2020) and orthologous COR47 (At1g20440.1), as well as the related COR78 (At5g52310.1), also showed relative increases in abundance after CA of *Arabidopsis* (Miki *et al*. 2019). It is thought that dehydrins, which are intrinsically disordered PM-associated proteins, act as chaperones to prevent protein denaturation during dehydration (Singh *et al*. 2019). These proteins likely interact with sucrose to change the glass transition temperature, decreasing the probability of ice crystal formation at sub-zero temperatures (Wolkers *et al*. 2001). Consistent with this finding, increasing levels of osmoprotectants including raffinose, glucose, and a 13-fold increase in sucrose were observed in CA2 plants (Figure 1). It should be recalled that under freeze conditions, most free water is bound to ice, thus a PM-dehydrin is likely critical to freezing tolerance and it is not surprising that the relative abundance would be increased to such a high level.

Other proteins that increased approximately two-fold in relative abundance by CA2 also appear to be important for PM protection. For example, a protein closely related to 97 kDa HSP (Bradi1g32770.1) could act similarly to COR410 and serve as a chaperone for PM stabilization. Phosphatidylinositol transfer protein (Bradi2g09760.1) ferries phospholipids to the PM and would also play a key role in PM restructuring; indeed the abundance of glycosylphosphatidylinositol-anchored proteins in *Arabidopsis* has been reported to increase almost two-fold after CA (Takahashi *et al*. 2016).

At CA2, 97 proteins showed a significant decrease in relative abundance (File S6). Notably, an estimated 42% of these proteins were annotated not as PM proteins but as cytoplasmic proteins. PM proteins can be reversibly associated with lipids or other proteins on the membrane (Marmagne *et al*. 2004). Thus it is possible that some of these proteins could be loosely associated with the PM but as low-temperature-induced restructuring of the PM commenced and continued, such associations could weaken or strengthen, resulting in their apparent decrease or increase in relative abundance, as has been reported for cold-stressed rice leaves (Komatsu *et al*. 2006). Proteins known to be associated with PMs under different conditions and that decreased in relative abundance were dominated by the so-called “master enzymes”, PM H^+^ ATPases (PM ATPase, Bradi5g24690.1; PM ATPase 1-like, Bradi1g54847.1) as well as other ATPases (V-type ATPase, Bradi1g67960.1; AAA+ ATPase, Bradi1g48010.1). Such PM ATPases are crucial for cell growth and maintain the transmembrane electrochemical gradient necessary for nutrient uptake (Morsomme and Boutry 2000). Thus, similar to other plants including *Arabidopsis*, there were relative abundance decreases in proteins annotated as having functions in cell growth, metabolism, and signal transduction (Huot *et al*. 2014; Fürtauer *et al*. 2019). Thus *Brachypodium* likely generally redirects energetics towards low-temperature protection, and away from growth as a general response during plant CA.

### The sustained response to cold acclimation

As CA progressed over several days, dynamic changes in the PM-associated proteome were observed. Heatmaps showed patterns consistent with longer CA (CA6 and CA8) showed similar PM-proteome patterns that were distinct from the shorter exposure patterns (CA2 and CA4), again suggesting an early low-temperature response followed by a sustained response. By CA6, 224 PM proteins were increased in abundance, 167 more than at the earlier time point. Of these, 20% (45) and 9% (21) showed fold-changes upwards of two or three orders of magnitude, respectively. The protein showing the greatest relative increase (∼55,000-fold) was an ABC transporter C-family member 5 like protein (Bradi1g75590.1). Such transporters have been previously reported to function in abiotic and biotic stress responses through cellular detoxification, and through compound exchange of hormones, metabolites, and defense molecules (Kang *et al*. 2011; Hwang *et al*. 2016). After prolonged low-temperatures, their role in detoxification is likely key to *Brachypodium* survival. Research with cold-stressed *Arabidopsis* has shown that similar ABC transporters have phosphorylation-level regulation within the membrane, further highlighting their importance (Kamal *et al*. 2020).

As noted, dehydrins interact with soluble carbohydrates, and after prolonged CA, sucrose reached concentrations in the leaves as high as 38 mg g^-1^. The time course of the increase in sugar concentrations in CA *Brachypodium* is similar to that reported for *Arabidopsis* as well as cereal crops (Plazek *et al*. 2003; Kamata and Uemura 2004; Klotke *et al*. 2004). Although this concentration is sufficient to lower the freezing point by a fraction of a degree, more importantly sucrose has additional roles in directly stabilizing membranes (Strauss and Hauser 1986) and can scavenge ROS even more efficiently than dedicated scavenging enzymes (Nishizawa-Yokoi *et al*. 2008; Stoyanova *et al*. 2011; Tarkowski and Van den Ende 2015). Sucrose is also involved in signaling and sugar synthesis activation (Kooiker *et al*. 2013), and the “sweet immunity” that aids in pathogen recognition (Duran-Flores and Heil 2016).

Coincident with the increase in sugar concentrations over the course of CA, a sugar transporter, (SWEET; Bradi2g24850.1), and a sucrose transporter SUT1-like protein (Bradi1g73170.1) increased in relative abundance, with two sucrose synthases (Bradi1g46670.1; Bradi1g60320.1) and two bidirectional SWEET sugar transporters (Bradi2g11920.1; Bradi2g56890.1) decreasing in relative abundance. Glycerol accumulation has also been shown to enhance abiotic stress resistance including cold resistance (Eastmond 2004). Although we did not assay for glycerol, we noted that glycerol kinase (Bradi5g23940.1) decreased in abundance after CA, suggesting reduced glycerol degradation allowing for accumulation of the polyol. After a week at low-temperature, *Arabidopsis* also showed relative increases in monosaccharide and sucrose transporters (Miki *et al*. 2019) indicating that soluble carbohydrate regulation is a common strategy for cold tolerance in evolutionarily distinct plants.

By CA6 there were 456 proteins displaying significant decreases in abundance (> 1.5-fold), approximately four-fold more than the number that decreased in abundance at CA2, including the PM ATPases (Bradi5g24690.1; Bradi1g54847.1; Bradi1g67960.1; Bradi1g48010.1). This observation is similar to reports from cold-stressed *Arabidopsis* (Muzi *et al*. 2016), but *Brachypodium* is notable for the large numbers of proteins that decreased in relative abundance during the sustained response, even with respect to the *Arabidopsis* dataset (Figure 2B,C). This again underpins our contention that normal growth in CA plants is arrested in order to utilize resources for stress responses, and *Brachypodium* may be more efficient than the dicot in this regard.

### Cross resistance to abiotic and biotic stresses

A network map enabled the visualization of relationships between the PM proteins whose abundances were influenced by CA, with its construct partially motivated by the observation that some other proteins could have roles in stresses that could accompany low-temperatures (Figure 5). Cold stress and pathogen stress represented the largest numbers of interactions (41 and 39, respectively) with proteins known to be associated with drought, osmotic, and salt stress clustering with the low-temperature responses, as might be expected from the known freeze-induced exclusion of solutes and cellular dehydration. The network indicates numerous instances of “crosstalk” among identified proteins, strongly suggesting that CA can “prime” *Brachypodium* for resistance to other abiotic and biotic stresses.

STRING was used to interrogate protein-protein interactions in the CA-induced PM proteome, which led to the identification of protein-interaction clusters associated with protein synthesis, and metabolism and energy production (Figures 6 and 7). The translational apparatus showed numerous interactions, but overall, there were hundreds of proteins with reduced relative abundance after CA that appeared to reflect a change in metabolism and the redirection of energy away from growth.

Translation reduction shown in cold-treated *Arabidopsis* triggers an intracellular increase in Ca^2+^ with the consequent regulation of cold-responsive genes (Guo *et al*. 2002; Zarka *et al*. 2003). Interactions in *Brachypodium* were generally conserved in CA2 and CA6, albeit with more interactions between the greater number of proteins identified after the longer cold exposure.

Cross resistance to a variety of stresses is evolutionarily adaptive since plants are rarely exposed to a single stress and extensive crosstalk exists between biotic and abiotic stress response pathways (Fujita *et al*. 2006; Rejeb *et al*. 2014). Subsequent to CA, immune signalling involved a host of HSPs and chaperones as well as peptidyl-prolyl cis-trans isomerases, a respiratory burst oxidase homolog protein B-like involved in pathogen resistance, and calcium-dependent protein kinase 13, as detailed in Results. This strongly suggests the activation of immune signaling pathways as a result of cold stress.

A single example of a protein with many interactions at CA6, allene oxide synthase 3 (Bradi3g08250.1), involved in the biosynthesis of jasmonic acid and plant defense (Farmer and Goossens 2019), showed 137 interactions with other proteins associated with functions in protein synthesis, as well as disease and defense. Thus the number and the identification of protein-protein interactions shows an obvious link between low-temperature stress and defense. The incidence and severity of bacterial and fungal diseases is known to be influenced by abiotic stresses including drought, extreme temperatures, or salinity (Moeder and Yoshioka; Snapp 1992; Koga *et al*. 2004; Freeman and Beattie 2009). With regards to low-temperature stress, wilting and necrosis caused by freezing damage can provide new entry points for pathogens, making plants susceptible at even low levels of infection. As well, psychrotolerant pathogens that thrive under the low ambient temperatures are more abundant, with some having the ability to initiate ice crystal formation at high sub-zero temperatures (Lindow *et al*. 1982; Wu *et al*. 2014). Indeed, CA winter cereals show higher disease resistance following low-temperature exposure (Kuwabara and Imai 2009; Wu *et al*. 2014; Pogány *et al*. 2016) and certain genes are similarly upregulated in response to both low-temperature and exposure to pathogens (Yeh *et al*. 2000; Wu *et al*. 2014). As such, plants that are tolerant to one stress are likely to have elevated tolerance to another stress as is evidenced by our meta-analysis. It is possible that such cross-resistance could be exploited for the production of crops that are better suited to multiple abiotic stresses associated with our changing environment.

### Conclusions

Taken together, we demonstrate the successful application of a two-phase partitioning system in combination with label-free quantification of proteins to achieve a PM enriched proteome in the model monocot, *Brachypodium*. In addition to identifying promising new protein and gene targets for research into cold tolerance, we have provided large scale datasets for future researchers to mine and analyze, and have importantly shown that changes in PM protein accumulation during days of CA are both dynamic and effective. We have also seen that monocots likely share certain freeze-resistance pathways and strategies with those that may be better known in dicots. Together our increased understanding of low-temperature responses represents an additional step towards enhanced cold tolerance in crops and a reduction in the sizable economic losses stemming from crop destruction due to increases in frost events attributed to our modern climate-mediated crisis.

## AUTHOR CONTRIBUTIONS

MB, TN, and HI conducted the experiments. CLJ, MB, TN, YK and MU analyzed the data. CLJ prepared the figures. CLJ, MB, GCD, and VKW wrote the paper, and all authors contributed to manuscript revision. MU, GCD, and VKW supervised the work. MU and VKW secured funding for the work.

## FUNDING

This research was funded by Kakenhi (#22120003 and #25650090) from the Japan Society for the Promotion of Science (JSPS) to MU and an NSERC (Canada) Discovery grant to VKW. MB was partially supported by a summer fellowship from the Japan Society for the Promotion of Science.

## CONFLICT OF INTEREST

The authors declare no conflicts of interest.

## DATA AVAILABILITY

All data described in this manuscript is original and produced by the authors. Supplemental files available at FigShare. File S1 contains all scripts and code used for the analysis of the *Brachypodium* cold-acclimated PM proteome. File S2 contains the stress response network meta-analysis for the CA2 dataset of the *Brachypodium* PM proteome. File S3 contains the total raw *Brachypodium* PM proteome dataset.

File S4 contains the significantly increased *Brachypodium* PM protein dataset. File S5 contains the significantly decreased *Brachypodium* PM protein dataset. File S6 contains the CA2 decreased annotated *Brachypodium* PM protein dataset. File S7 contains the CA6 decreased annotated *Brachypodium* PM protein dataset. File S8 contains the CA2 increased annotated *Brachypodium* PM protein dataset. File S9 contains the CA6 increased annotated *Brachypodium* PM protein dataset. File S10 contains the interactive network for the stress response meta-analysis of *Brachypodium* CA2 proteins. File S11 contains the interactive network for the CA2 predicted protein-protein interactions. File S12 contains the interactive network for the CA6 predicted protein-protein interactions. File S13 contains the CA2 preliminary *Brachypodium* PM dataset.

## Supplemental materials

**Figure S1.**
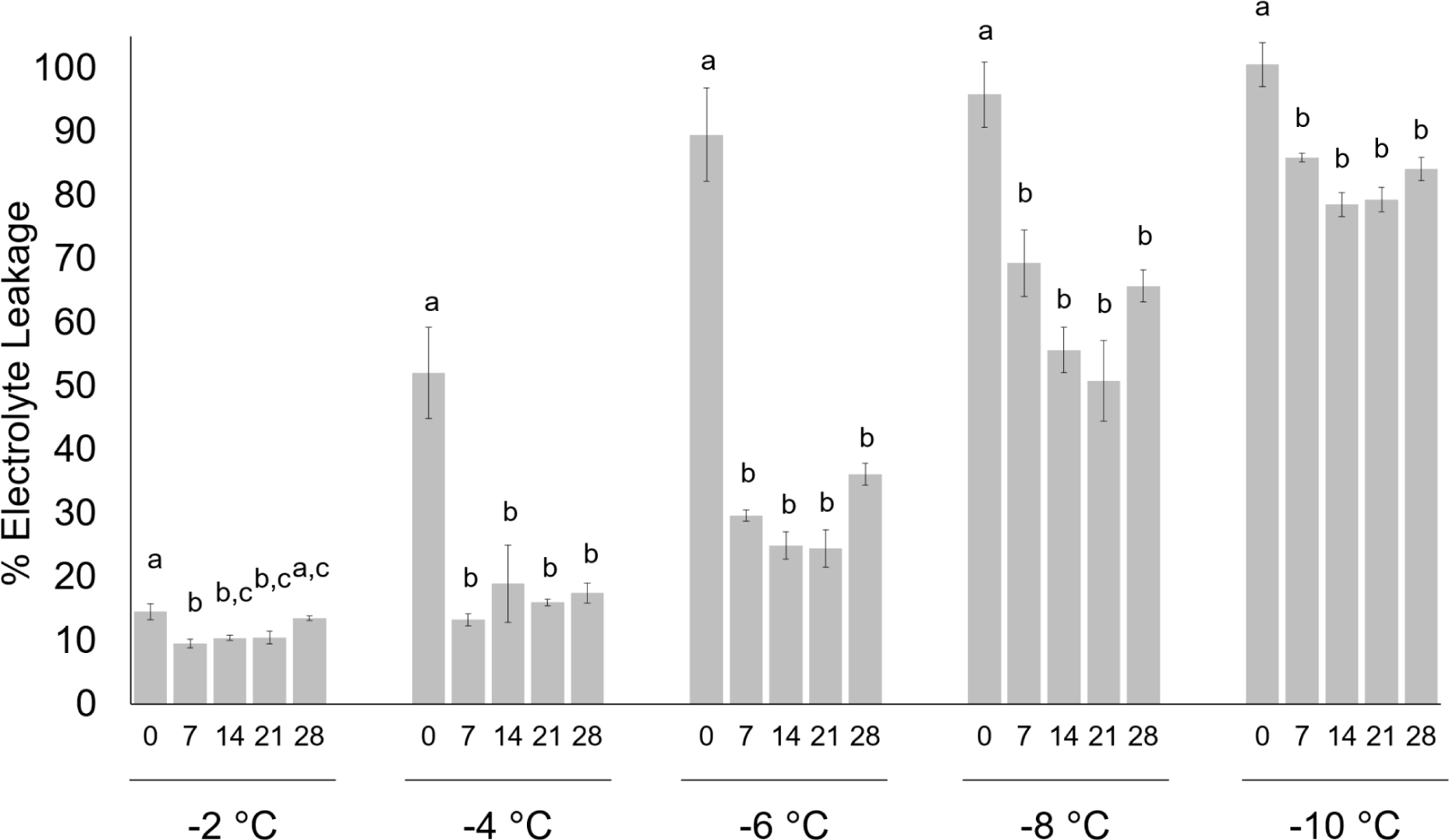
Electrolyte leakage assays conducted using leaf tissue from non-acclimated and cold-acclimated wildtype *Brachypodium distachyon* ecotype *Bd*21. Days of cold acclimation and temperatures are shown. Samples were frozen to decreasing final temperatures at a rate of 1 °C every 15 min before being assayed for electrolyte leakage (%). Four biological replicates were conducted (n = 10) and ANOVA and post-hoc Tukey tests were performed. Error bars represent standard error of the mean. Letters indicate statistically significant groups (*p* < 0.05) with analyses conducted on separate groups for each temperature.

**Figure S2.**
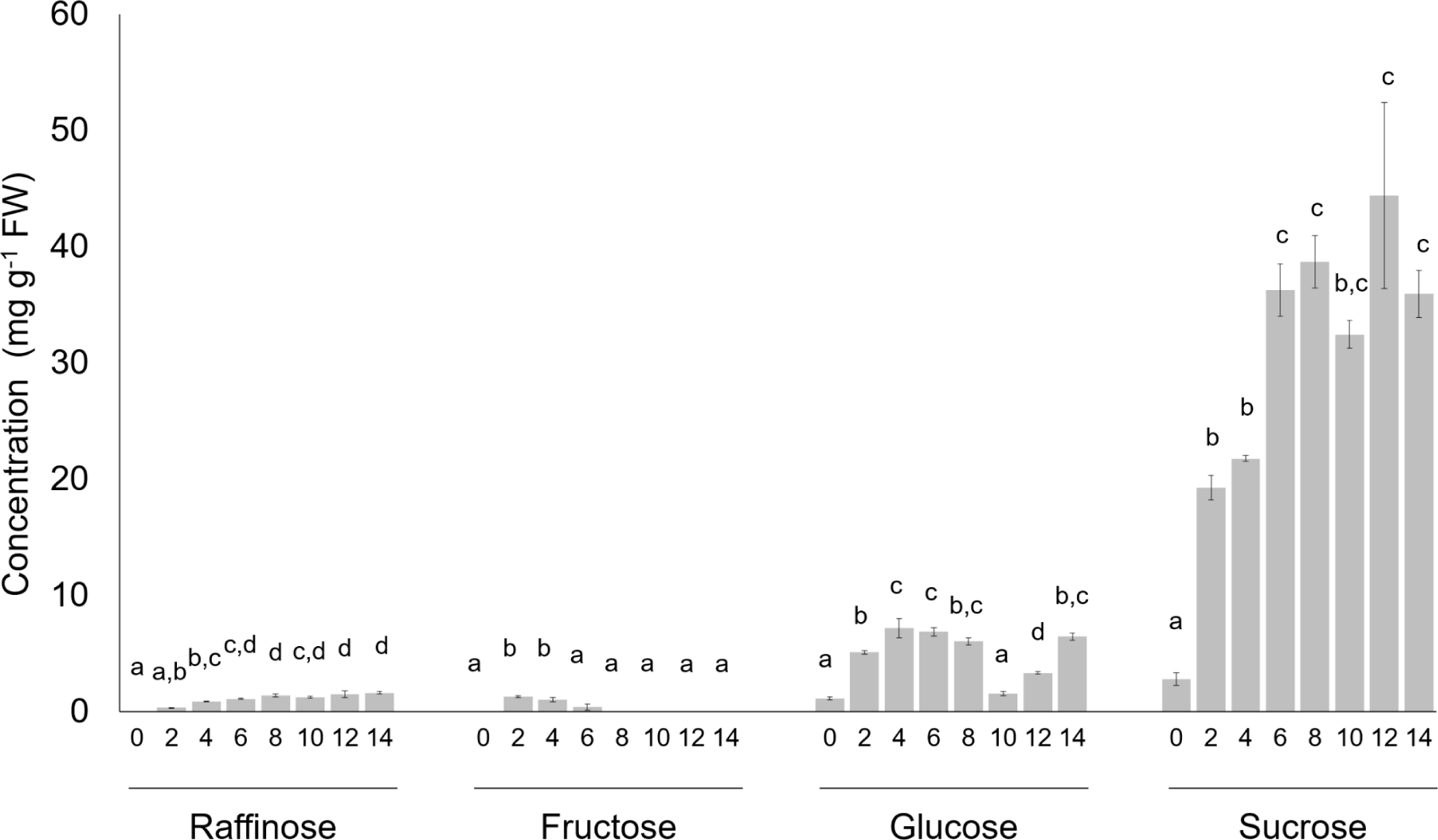
Accumulation of soluble sugars in leaf tissue from non-acclimated and cold-acclimated wildtype *Brachypodium distachyon* ecotype *Bd*21. Days of cold acclimation and soluble sugar concentration in fresh weight (FW) of leaves as shown. Four biological replicates were conducted (n = 10) and ANOVA and post-hoc Tukey tests were performed. Error bars represent standard error of the mean. Letters indicate statistically significant groups (*p* < 0.05) with separate analyses conducted for each soluble sugar.

**Figure S3.**
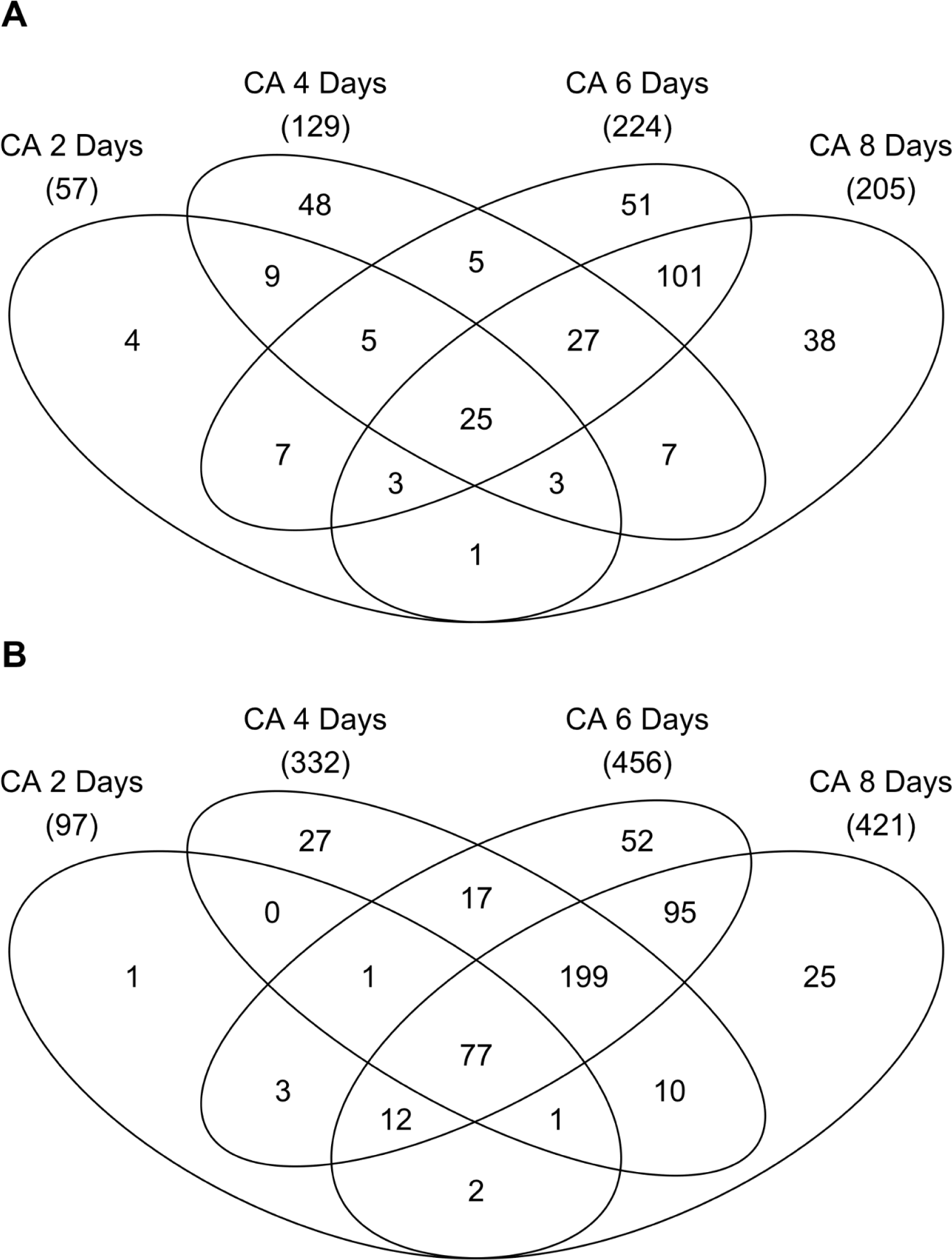
Venn diagrams of the total number of *Brachypodium distachyon* proteins with significant differences in relative accumulation over the entire low-temperature time course. (A) Proteins that increased in abundance with fold-changes > 1.5 and **(B)** proteins that decreased in abundance with absolute fold-changes > 1.5 following two, four, six, and eight days of cold acclimation (CA).

**Figure S4.**
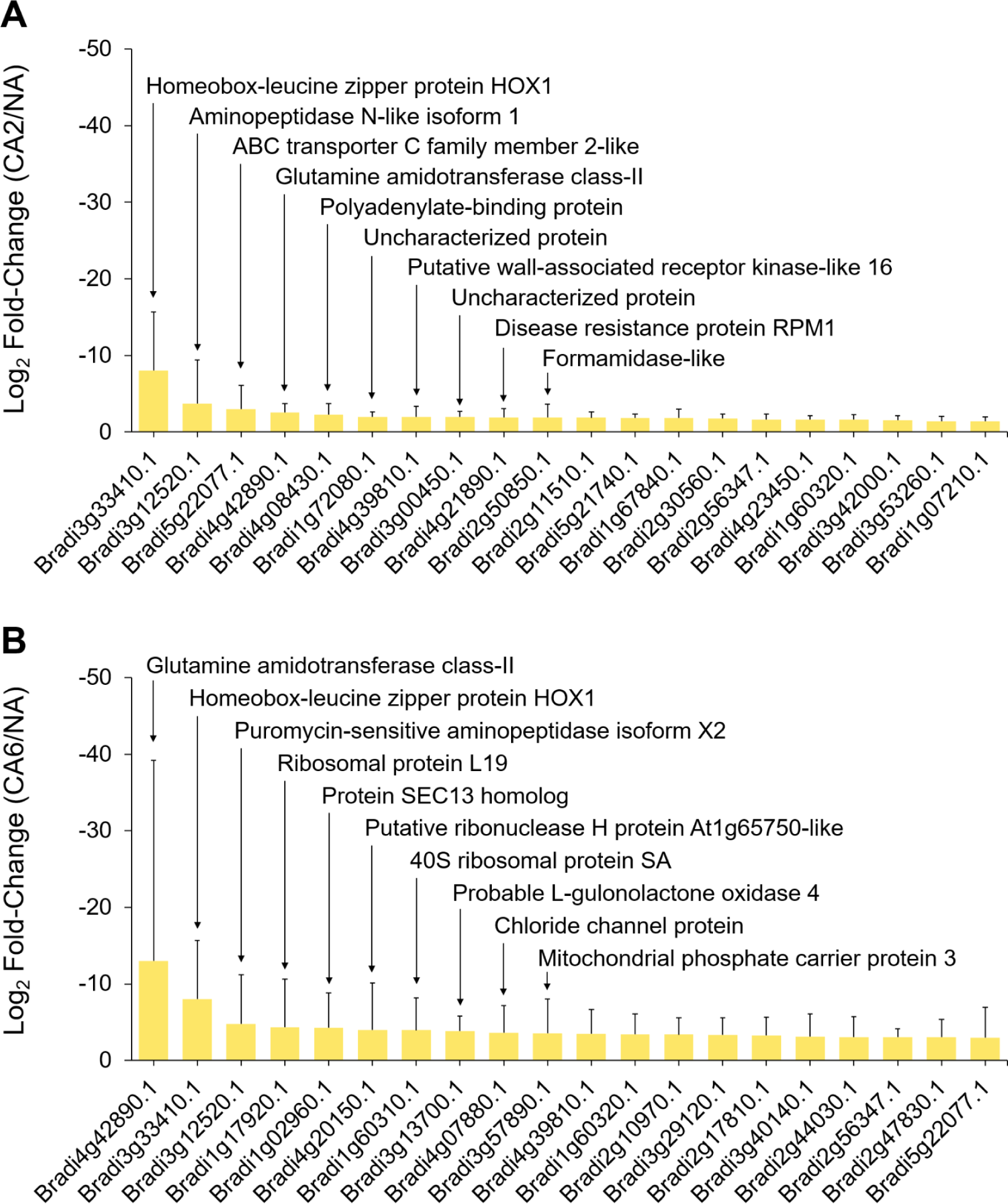
*Brachypodium distachyon* proteins undergoing the greatest fold decreases in relative abundance during cold acclimation. Data is shown for **(A)** two days and **(B)** six days of cold acclimation. Values are the average of four replicate trials and protein annotations were predicted as previously described. Error bars represent standard error of the mean. A pseudo-count of one was added to the cold-acclimated value for proteins that were not detected under that experimental condition. The ten proteins with the greatest fold relative increases are labelled accordingly. A log _10_ scale is shown.

